# Integrated integral population models

**DOI:** 10.1101/2023.10.24.563870

**Authors:** Paola Portillo-Tzompa, Paulina R. Martín-Cornejo, Edgar J. González

**Affiliations:** Departamento de Ecología y Recursos Naturales, Facultad de Ciencias, Universidad Nacional Autónoma de México. Circuito Exterior s/n, Ciudad Universitaria, 04510, Mexico City, Mexico; Posgrado en Ciencias Biológicas, Universidad Nacional Autónoma de México. Circuito de Posgrados s/n, Ciudad Universitaria, 04510, Mexico City, Mexico

**Keywords:** Bayesian model, data integration, integral projection model, integrated population model, inverse estimation, matrix population model, Soay sheep, structured population dynamics

## Abstract

1. Data integration allows obtaining better descriptions and forecasting of a population’s behaviour by incorporating data at both the individual and population levels. Structured population models include the matrix population models (MPMs), which structure a population through a discrete state variable, and the integral population models (IPMs), which use a continuous variable. Two decades ago, the integrated version of MPMs appeared, but their corresponding version for IPMs is still missing.
2. Here, we propose the integrated integral population model (IIPM). This model takes up the ideas behind existing models used to describe and forecast the dynamics of continuously structured populations: IPMs, which use individual data, and inverse IPMs, which use population data. Particularly, we emphasise the construction and fitting of the IIPM under a Bayesian framework and use the Soay sheep database to compare the population dynamics generated by the IIPM and these existing models.
3. The IIPM constructed with the Soay sheep data had a good performance both at the individual (vital rates) and population (size and structure) levels, because, as they are constrained to fit both sets of data, they produce a balanced population dynamics. In turn, the IPM produced the best individual estimates and the worst population estimates, whilst the inverse IPM produced the worst individual estimates and the best population estimates.
4. The objective of a structured population model should be to correctly describe population patterns. IPMs, by not using population data, fail in this objective. An IIPM solves this problem.

## Introduction

The history of the study of population dynamics has been the history of the development of increasingly complex models (Benton et al., 2006; Metcalf & Pavard 2006; Evans, 2012; Riecke et al., 2019). These models aim to make a more detailed description of the structure of a population and the drivers that determine its dynamics (Ellner & Rees 2006; Schaub & Abadi, 2011; Ellner et al. 2016; Plard et al., 2019a; 2019b) This complexity goes hand in hand with a need for higher maths literacy among demographers, and computational power (Besbeas & Morgan, 2019; Plard et al., 2019b; Fung et al., 2022; Frost et al., 2023). In recent years, data integration has allowed us to obtain better descriptions and forecasting of a population’s behaviour by incorporating data at both the individual and population levels (Evans et al., 2016; Zipkin & Saunders, 2018; Zipkin et al., 2019; Frost et al., 2023).

Currently, data integration exists for population models where the population is structured by discrete variables, but their correlate for continuous variables is still missing (Schaub & Abadi, 2011; Zipkin et al., 2019; Frost et al., 2023). A revision of existing models allows for a clear overview of how this model comes about and can be readily constructed.

### Structured population models

Two popular statistical frameworks for modelling structured populations are matrix projection models (MPM) and integral projection models (IPM). Both models have the same goal, which is the description of the dynamics of a population, and the projection of its behaviour over time (Merow et al., 2014; Salguero-Gómez et al., 2015; 2016). However, they differ in the identity of the variable used to structure the population. While MPMs structure the population around a discrete variable, e.g., stage and sex, IPMs structure the population around a continuous one, e.g., size and age (Caswell, 2001; Ellner et al., 2016). In relating a variable with the vital rates determining the life cycle of an individual, i.e., its survival, growth and fecundity, regressions are a commonly used tool (Merow et al., 2014; Ellner et al., 2016). IPMs were the first in implementing regressions in the construction of the function describing the dynamics of a population, known as the kernel (Easterling et al., 2000). MPMs have commonly used simple estimates based on category-specific survey data and have just recently started to recognize the benefits of using regression in obtaining such estimates (Jäkäläniemi et al., 2011; Gremer et al., 2012; Alahuhta et al., 2017; Ramula et al., 2020). These regressions can be performed either within a likelihood (Ellner et al., 2016; Ramula et al., 2020) or a Bayesian framework (Ellner et al., 2016; Elderd & Miller 2016). In using regression, the main benefit is that estimates are based on the entire vital-rate data set instead of splitting it into categories (Ellner et al., 2016). However, these models, MPMs and IPMs, are fed only by individual level data (Fig. 1, Easterling et al., 2000; Caswell, 2001).

**Figure 1.**
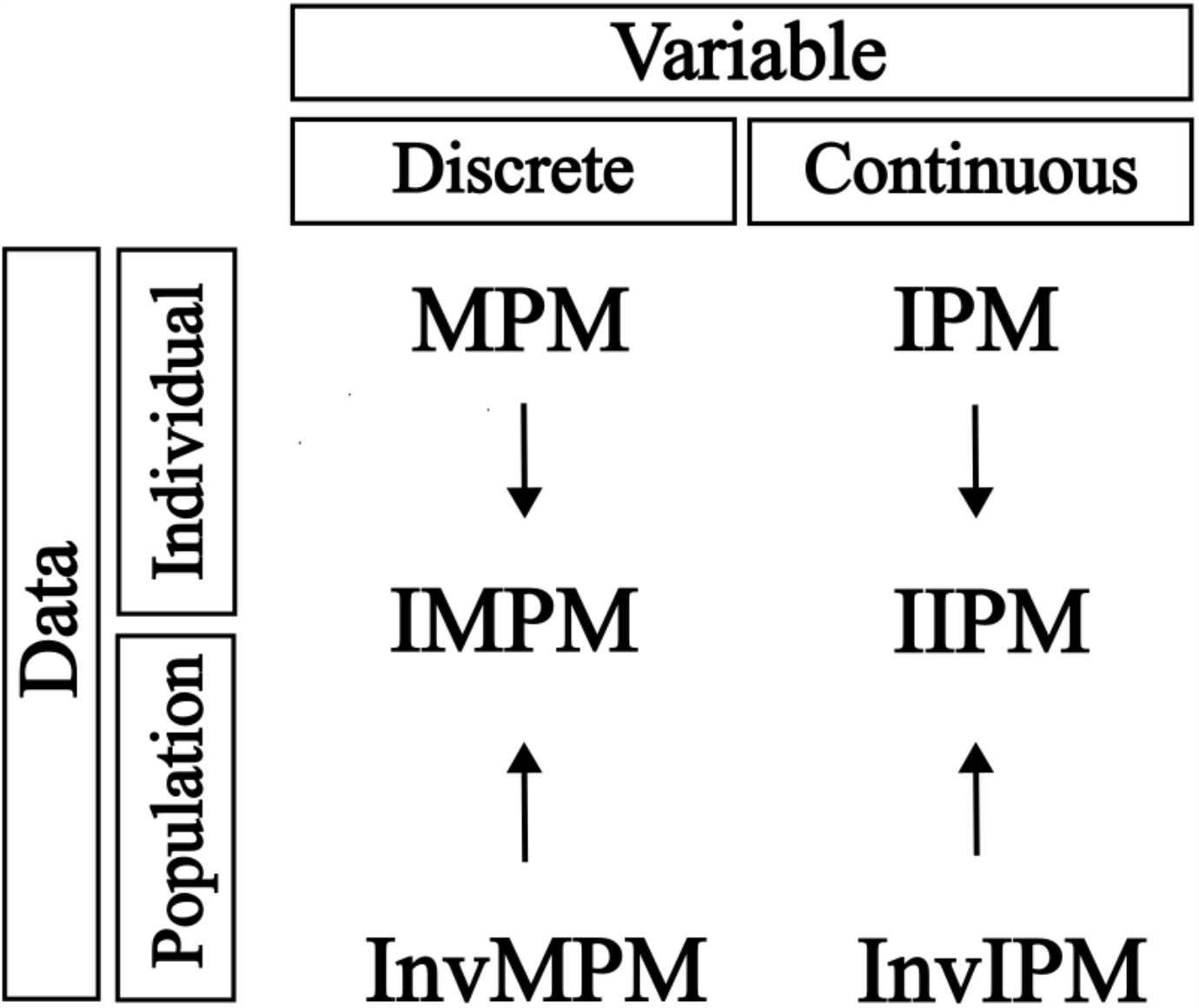
Location of the integrated integral population model with respect to the existing ones according to the type of data they use and the type of state variable that structures the population. MPM: Matrix Population Model, IMPM: Integrated (Matrix) Population Model, InvMPM: Inverse Matrix Population Model, IPM: Integral (Population) Projection Model, IIPM: Integrated Integral Population Model, InvIPM: Inverse Integral (Population) Projection Population Model. In parenthesis are the terms that we propose to unify the nomenclature of these models. Highlighted are the models that we compare in this study as they are more closely related with the IIPM.

As stated, the main goal of population models is to describe and project population attributes, such as the population size, and, in structured population models, the population structure (Caswell, 2001; Ellner et al., 2016). Thus, there is a mismatch between the level of input, i.e., individual data, and the level of the output, i.e., population attributes. This mismatch causes a lack of fit at the population level (Ghosh et al., 2012), because the fitted models are describing the dynamics of an average individual instead of the dynamics of the whole population. To handle this mismatch, we can model population data, like time series of population size and population structure, in different ways (Ghosh et al., 2012; Plard et al 2019a; 2019b). The first way consists of reconstructing the entire population dynamics through a model based only on population data; these are known as inverse population models (Fig. 1; Gross et al., 2002; Ghosh et al., 2012; González et al., 2016; Tredennick et al., 2017). The second way consists of doing this reconstruction based on both individual and population data (Plard et al 2019a; 2019b). We now describe both approaches.

### Inverse structured population models

Inverse population models contribute to the ecologist’s toolbox recognising the importance of population level data in population dynamics models (Gross et al., 2002; Ghosh et al., 2012; González et al., 2016). These models conceive the demographic core as a deterministic function that redistributes the population structure over time. Thus, variation in population time series reflects some of the demographic behaviour. In particular, inverse IPMs implement regressions for population time series, which can also be fitted within a likelihood (González et al., 2016) or Bayesian framework (Gross et al., 2002; 2012). However, because there is no individual data in inverse IPMs, their success is only achieved under very particular circumstances: in a likelihood framework, simple life cycle species with long population size time series, and size distributions composed by lots of individuals are needed (González et al., 2016); in a Bayesian framework, inverse models require very strong priors, so if not enough data are available, the posterior distributions of each parameter will be close to its prior (Gross et al., 2002; 2012). The importance of both individual and population data explains the problems associated with inverse population models and the emergence of the second way of fitting population dynamics models.

### Integrated (matrix) population models

Integrated population models combine individual and population data to describe the population dynamics through an MPM (Fig. 1, Shaub & Abadi, 2007; Zipkin & Saunders, 2018). Integration is a new trend in population ecology which recognises the benefits of incorporating multilevel data in a single population dynamics model (Evans et al., 2016, Zipkin & Saunders, 2018; Frost et al., 2023). This approach allows us to obtain better descriptions and forecasting of the population behaviour and solves the input-output mismatch of non-integrated models (Kéry & Shaub, 2011; Plard et al., 2019a; 2019b).

Notably, integrated population models, as they use an MPM at its core, are in fact integrated *matrix* population models (IMPMs) and are mainly used in the analysis of animal populations (Shaub & Abadi, 2011; Bled et al., 2017; Plard et al., 2019b; Gamelon et al., 2021; Moeller et al., 2021) due to the particular difficulties associated with the surveying of non-sessile organisms. These difficulties translate into having multiple methods to measure vital and population attributes, including capture-mark-recapture, telemetry, and non-invasive DNA data while population attributes have been modelled using aerial population estimates and population count data (Shaub & Abadi, 2011; Bled et al., 2017; Plard et al., 2019b; Gamelon et al., 2021; Moeller et al., 2021). The ability of IMPMs to jointly analyse the different databases associated with these methods explains their assimilation among the animal demographers (Shaub & Abadi, 2011; Zipkin & Saunders, 2018).

However, data integration is not exclusive to animal studies (Evans et al., 2016; Isaac et al., 2019). In plant population studies, the database could come from a single population surveyed over time (Merow et al., 2014). In this case, data integration is also important as we have information of two different biological levels affecting the same process (Isaac et al., 2019). Presently, data integration only exists for population models where the population is structured by discrete variables, i.e., where a matrix is an adequate tool (Schaub & Abadi, 2011; Plard et al., 2019b).

Here, we propose the integrated integral population model (IIPM; Fig. 1). This model takes up the ideas behind an IPM, InvIPM and IMPM to describe and forecast the dynamics of populations structured around a continuous state variable based on both individual and population data. Particularly, we emphasise the construction and fitting of the IIPM under a Bayesian framework using the Stan software (Stan Development Team, 2023). We use the Soay sheep database to compare the population dynamics generated by an IPM, InvIPM, IMPM and IIPM.

### Existing Integral Projection Models (IPMs)

IPMs model a population’s dynamics through a set of structured vital rates based on individual data (Fig. 1, Easterling et al., 2000; Ellner et al., 2016). A traditional IPM is defined as a recursive function

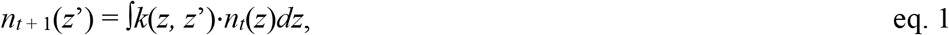

eq. 1 where *k*(*z, z*’) is the demographic deterministic function, called the kernel, describing the dynamics (Ellner et al., 2016) and *n*_*t*_(*z*) is the population structure, *z* is the state variable at time *t* and *z*’ is this variable at time *t* + 1. The population size, *N*_*t* + 1_, can be estimated by integrating *n*_*t* + 1_(*z*’) over *z*’, i.e.

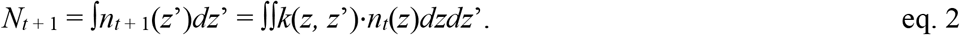

Henceforth, we will use individual size as the state variable.

The kernel is composed of vital rates functions describing the life cycle of the study species. A general structure of the kernel is

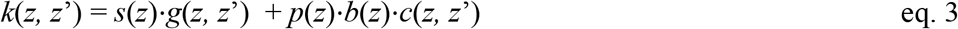

where *s*(*z*) is the survival probability of an individual of size *z, g*(*z, z*’) is the probability of an individual having a size *z* at time *t* of having a size *z*’ at time *t* + 1, *p*(*z*) is the probability of an individual of size *z* of reproducing, *b*(*z*) is the mean number of newborns that an individual of size *z* contributes to the population at the time *t* + 1, and *c*(*z, z*’) is the probability of a newborn produced by a size-*z* individual of having a size *z*’.

### IPM that uses individual data

To inform the IPM using individual survey data, we use information to estimate the parameters determining the structure of the vital rates functions (eq. 3). A traditional IPM consists of linear functions relating each vital rate with the state variable (*z*; Ellner et al., 2016). To estimate the parameters associated with the vital rates functions we use generalised linear models (GLMs). The survival and reproduction probability functions, *s*(*z*) and *p*(*z*), are the probabilities estimated through binomial GLMs, i.e.

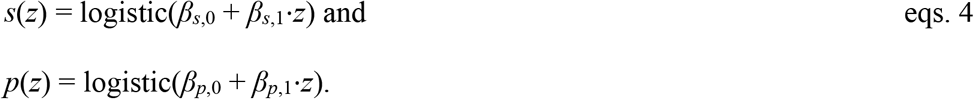

The growth and newborn sizes probabilities, *g*(*z, z*’) and *c*(*z, z*’), are Gaussian GLMs, i.e.

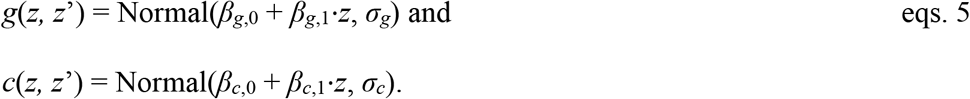

Finally, the function describing the mean number of newborns is the estimated mean of a negative binomial GLM, i.e.

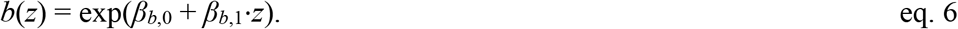

Under a maximum likelihood approach, we would estimate the set of parameters ***β*** and ***σ*** using data on the vital rates; through a Bayesian approach, we would estimate the posterior distributions of this set of parameters. In both cases, we need to calculate the likelihoods associated with the vital rates, i.e., *L*(**s** | ***β***_*s*_), *L*(**g** | ***β***_*g*_,*σ*_*g*_), *L*(**p** | ***β***_*p*_), *L*(**b** | ***β***_*b*_,*φ*_*b*_) and *L*(**c** | ***β***_*c*_,*σ*_*c*_), for *s*(*z*), *g*(*z, z*’), *p*(*z*), *b*(*z*) and *c*(*z, z*’), respectively; where ***β*** are the parameters associated with the mean of the distribution and *σ* to its dispersion, with subscripts indicating particular vital rates. Note that, here and throughout the rest of the text, we use light typeface for variables and functions (e.g., *s, z, N*_0_), lowercase bold for vectors (e.g., **s, *β***), and uppercase bold for matrices; and we use regular typeface for data (e.g., **s**) and italics for estimated parameters (e.g., *σ*_*g*_, ***β***).

Finally, the likelihood of the entire IPM using individual data is the multiplication of the likelihoods associated with the vital rates, i.e.,

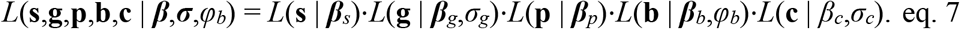

### IPM that uses population data: Inverse Integral Projection Model (InvIPM)

InvIPMs are fed only by population data and estimate the parameters associated with the vital rates at the population level (Gosh et al., 2012; González et al., 2016). Thus, InvIPMs reconstruct the kernel (eq. 3), and they have the advantage that, given that they use population data, they produce a better description of the observed size structures, **n**_*t* + 1_, compared with an IPM (González et al., 2016). Furthermore, to iterate the kernel function InvIPMs start with an estimated initial size probability distribution, *d*_0_(*z*), and an estimated initial population size, *N*_0_, such that *n*_0_(*z*) = *N*_0_·*d*_0_(*z*).

Thus, an InvIPM is defined through two GLMs. For the population structures, we fit a GLM with an empirical distribution. This distribution corresponds to the size probability distribution derived from the IPM iteration (eq. 1), i.e.,

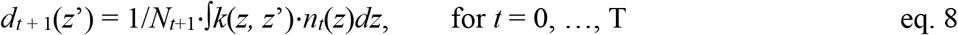

where *N*_*t*+1_ (as calculated through eq. 2) serves as a proportionality coefficient to make *d*_*t*+1_(*z*’) integrate to 1.

For the population sizes, a second negative binomial GLM allows to estimate the mean population size at each time, i.e.,

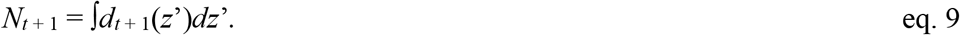

Multiplying the likelihoods associated to these two GLMs, we get the likelihood of the entire InvIPM as

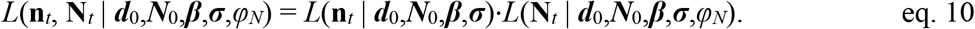

### MPM that uses individual and population data: Integrated Matrix Population Model (IMPM)

IMPMs are MPMs that are fed by individual and population data (Fig. 1, Besbeas et al., 2002; Schaub & Abadi, 2011). As with all MPMs, the population is structured through a discrete state variable; here we assume that a simple two-stage age variable (juveniles and adults) structures the population. The individual data consist of the survival (**s**_j_ and **s**_a_), reproduction (**p**_j_ and **p**_a_), and number of newborns produced (**b**_j_ and **b**_a_) by an individual at each stage. The population data used is a time series of size structures **n**_*t*_ = [n_j,*t*_, n_a,*t*_], such that N_*t*_ = n_j,*t*_ + n_a,*t*_. With this information, we seek to estimate, at the individual level, the survival probabilities (*s*_j_ and *s*_a_), the reproduction probabilities (*p*_j_ and *p*_a_), and the average number of newborns produced (*b*_j_ and *b*_a_) by an individual at each stage; and, at the population level, the initial proportions of juvenile and adult individuals (***n***_0_ = [*j*_0_, *a*_0_]) and the initial population size (*N*_0_). Thus, the demographic core of the IMPM, equivalent to the kernel in an IPM, is a 2 × 2 transition matrix

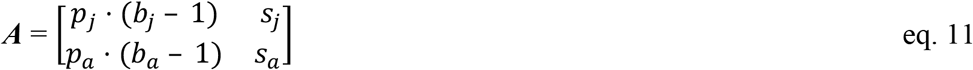

The iteration of this matrix, starting with ***n***_0_, produce a time series of estimated size structures through

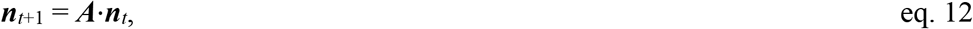

which is equivalent to eq. 1. This can also be expressed as

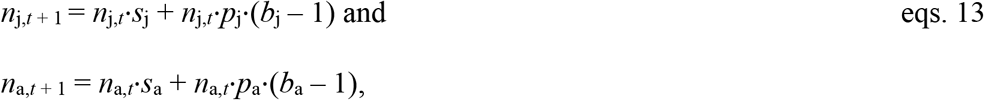

a more traditional way to express IMPMs.

The binomial GLMs for the vital rates are defined as

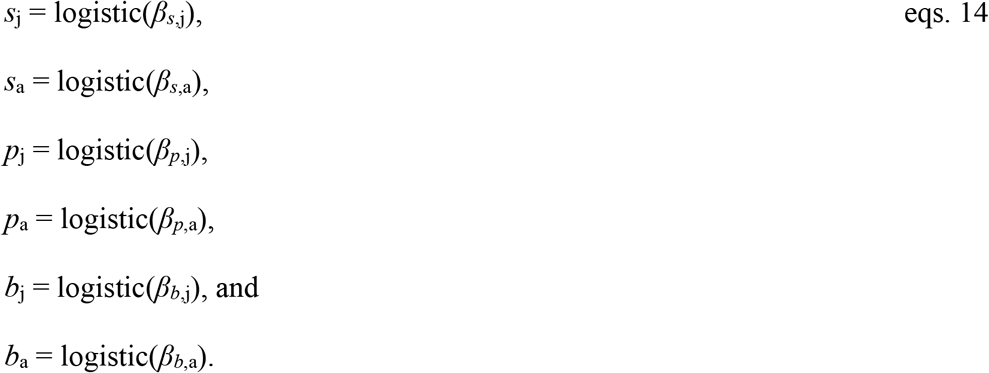

which are equivalent to eqs. 4–6.

At the population level, the GLMs for the population data are two GLMs. For the population structures, we fit a binomial GLM with mean probabilities

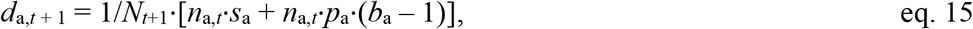

following eqs. 13. Note that we use a binomial distribution because the population is structured by a binary variable; if the population were structured by a variable with more than two categories a multinomial distribution would be required.

In turn, for the population sizes, we fit a negative binomial GLM with mean

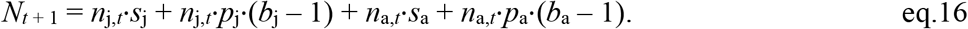

Thus, the likelihood associated to an IMPM is

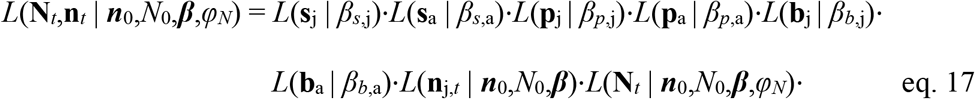

### How to build an Integrated Integral Population Model (IIPM)

IIPMs combine IPMs and InvIPMs. Thus, IIPMs are fed by both individual and population data (Fig. 1). With individual data we estimate the vital rates parameters, using eqs. 4-6. With population data we estimate the initial size structure, ***n***_0_(*z*’), and the initial population size, ***N***_0_, using eqs. 8-9. Therefore, the likelihood associated to an IIPM is

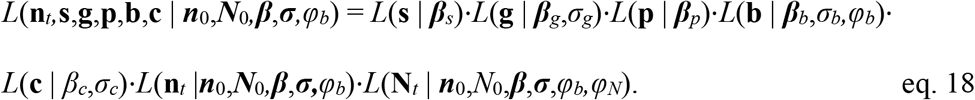

### An application to a simple study system: Soay sheep

To apply IPMs, InvIPMs, IMPMs and IIPMs we used the well-known Soay sheep (*Ovis aries*) database. This database is composed of demographic data of female sheep surveyed in Village Bay, Hirta Island, Scotland, UK, between 1986 and 1996 (Clutton-Brock & Pemberton, 2004; Coulson, 2012). We built the IPM, InvIPM, and IIPM using individual weight (kg) as the state variable (*z*). For the IMPM we used the juvenile (≤ 1 yr old) vs adult (> 1 yr old) dichotomy as the state variable. Also, we took into account that a female can produce a maximum of two newborns per year. Thus, we modelled the number of newborns as a binomial GLM, i.e.,

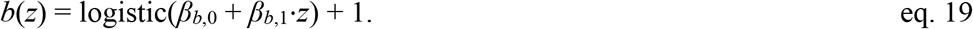

### Settings

For the IPM, InvIPM, and IIPM, their numerical implementation used a kernel partition of 50. We constructed all models in the Stan software (Stan Development Team, 2023).

We used the default prior distributions set by the brms package (Bürkner, 2021). For model diagnosis, we checked the R-hat to be 1 and the effective sample size (ESS) to be over 1000 for all parameters in all models. Finally, to compare between models, we estimated, for each model, the pointwise log-likelihood of the data of each vital rate and population attribute; the model with the largest pointwise log-likelihood has a better fit to the data (Vehtari et al., 2017).

## Results

The IIPM for the Soay sheep data estimated a population dynamics that produces good fits both at the individual and population levels.

At the individual level, as expected, the IPM had the best performance (in terms of the pointwise log-likelihood), among the three IPMs being compared (Table 1); this because the IPM’s only aim is to fit the individual level data; the IIPM had a close performance in the estimation of vital rates parameters to that of the IPM; and the InvIPM had the worst performance (Table 1, Fig. 2). Comparing the IMPM with the IIPM, the former had a performance worse than the latter (Table 1).

**Table 1.**
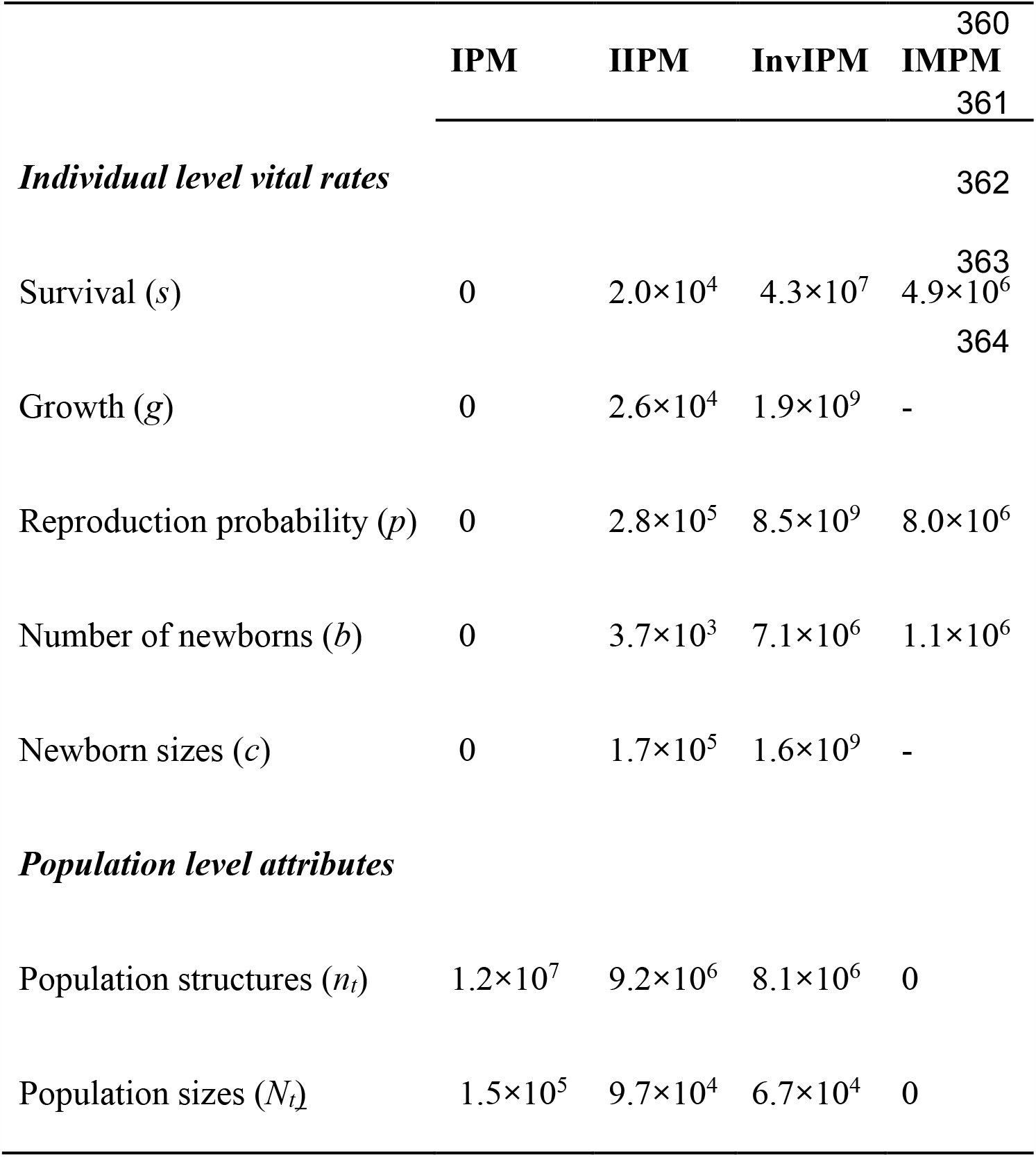
Pointwise log-likelihoods for the five vital rates and two population traits obtained for the four population models compared in this study: IIPM: Integrated Integral Population Model, IPM: Integral Population Model (IPM), Inverse Integral Population Model (InvIPM), IMPM: Integrated Matrix Population Model. Since the IMPM is not structured by size it does not have a log-likelihood associated with the growth (*g*) and newborn size distribution (*c*) functions.

**Figure 2.**
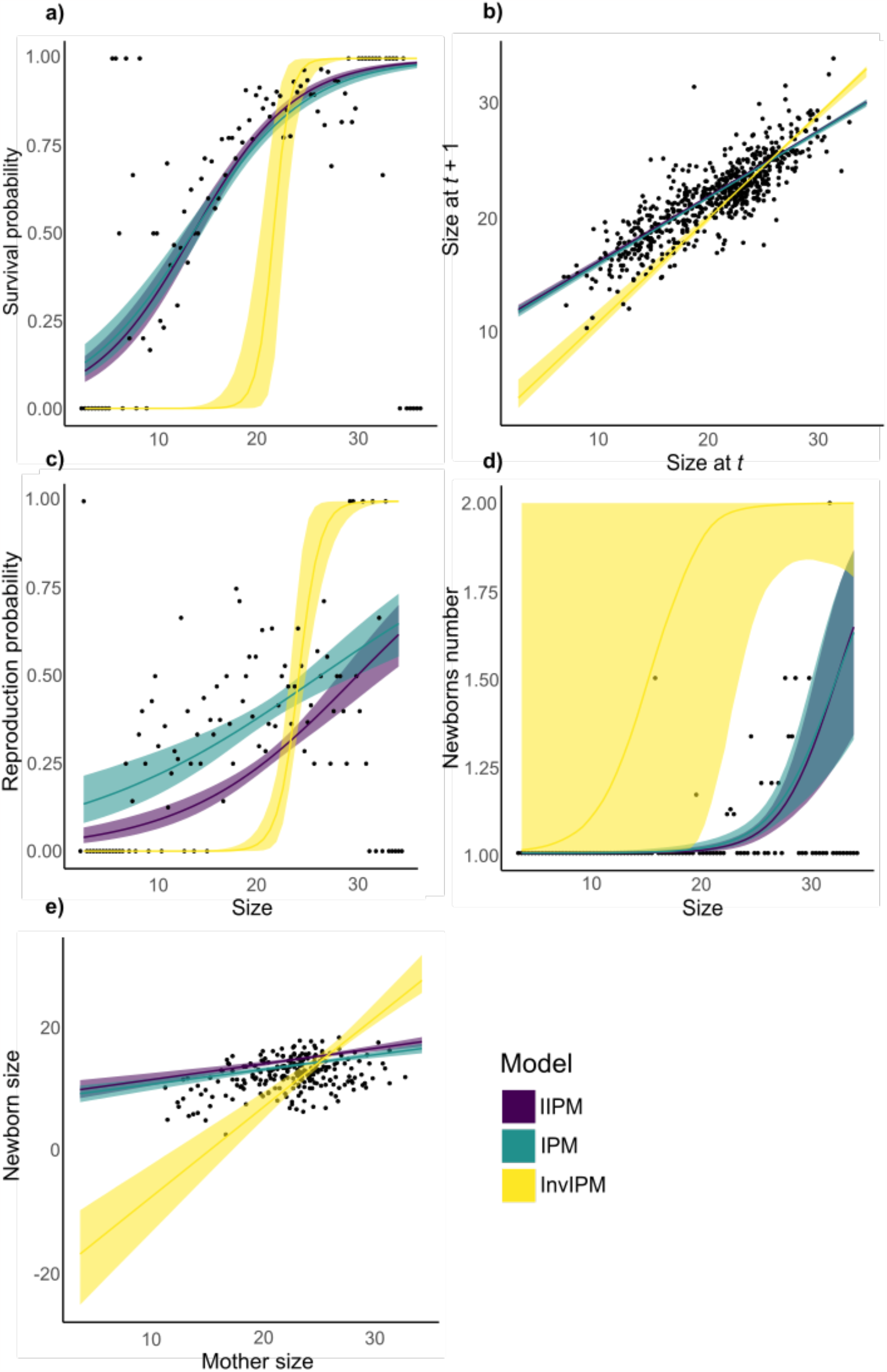
Predicted patterns for the five vital rates that jointly determine the dynamics of the Soay sheep population according to the three population models structured by a continuous variable (weight, kg): IIPM: Integrated Integral Population Model, IPM: Integral Population Model (IPM) and Inverse Integral Population Model (InvIPM). We include the 95% credibility bands associated with each predicted pattern. Individual data are in black.

In comparing the IIPM with the best performing model, the IPM, we see that they predicted similar patterns in the relationship between the state variable and the vital rates (Fig. 2). The survival probability of sheep increased smoothly with weight in both models (Fig. 2a); they also predicted a mean growth that is in agreement with the data, with lightweight sheep increasing their size, while heavy ones decreasing it (Fig. 2b). Both models predicted that the probability of reproduction increases with size, with the IIPM estimating a similar trend than that of the IPM, but with lower reproduction probabilities for almost all sizes (Fig. 2c). Additionally, these models predicted that the mean number of newborns produced in a single reproductive event increases with weight; on average, lightweight sheep give birth to one sheep and heavy ones to twins (Fig. 2d). Finally, both the IIPM and the IPM predicted that heavier mothers produce heavier newborns (Fig. 2e).

At the population level, as expected, the InvIPM had the best performance among the three IPMs being compared but, interestingly, the IMPM performed even better (Table 1). This result is probably related to the structure of the matrix of the IMPM. This model structured the population using age as the state variable, while all the continuous models structured the population using weight. This suggests that, for the Soay sheep population, age is a better explanatory state variable than weight. Thus, it is the task of the researcher to identify the state variable that best describes the population dynamics.

All IPM models predicted that the size distribution fluctuated in the first years and then stabilised (Fig. 3); however, the IPM underestimated the proportion of heavy individuals and overestimated the proportion of medium weight individuals in almost all years. In turn, all models clearly identified the increase over time in the observed population size (Fig. 4); however, the IIPM, IPM and IMPM identified an acceleration of this increase, whilst the InvIPM identified a deceleration.

**Figure 3.**
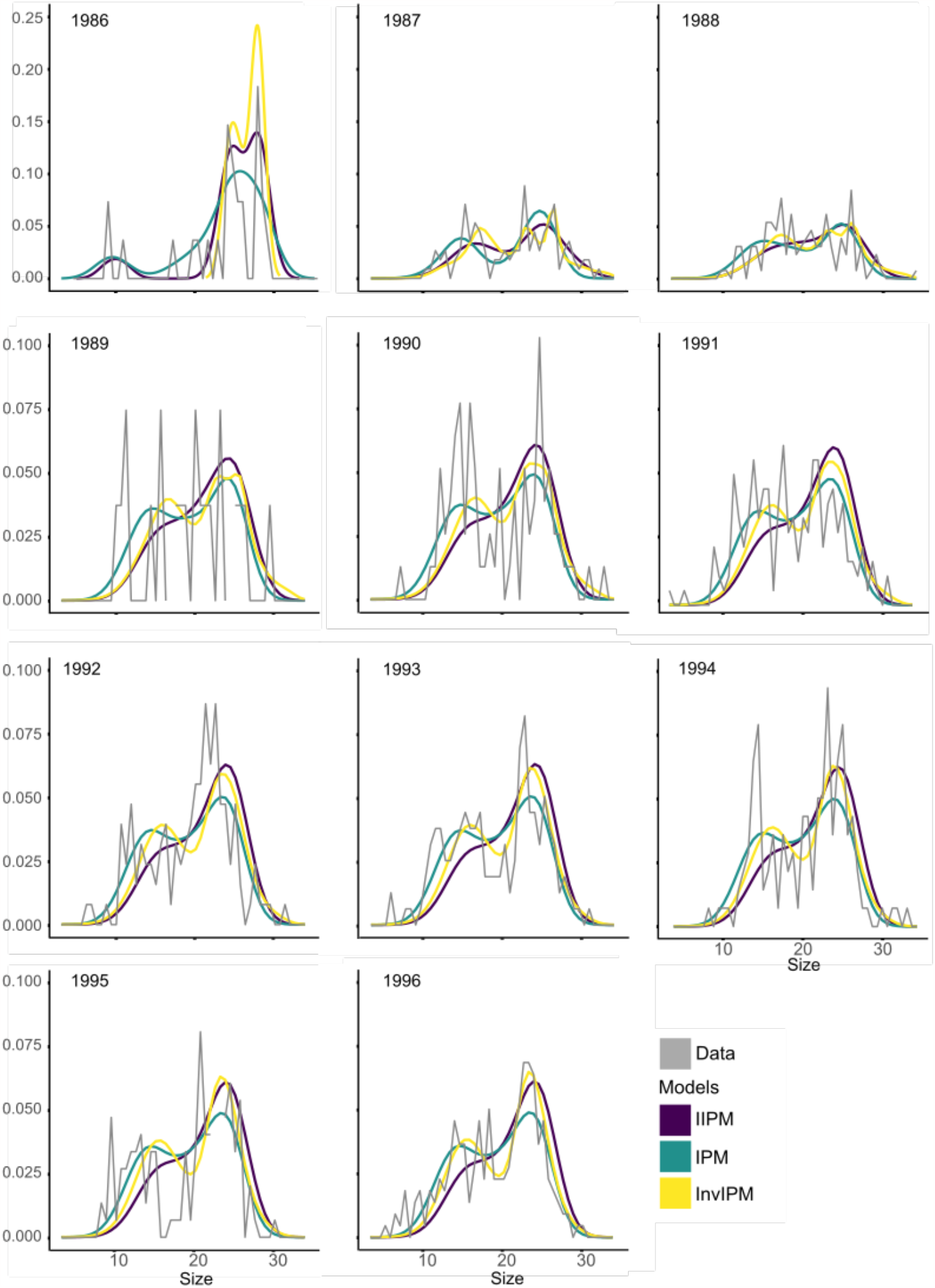
Predicted size structures of the Soay sheep population over 11 years according to the three population models structured by a continuous variable (weight, kg): IIPM: Integrated Integral Population Model, IPM: Integral Population Model (IPM) and Inverse Integral Population Model (InvIPM). Observed population size structures are in grey. Note that the y-axis scale changes after 1988.

**Figure 4.**
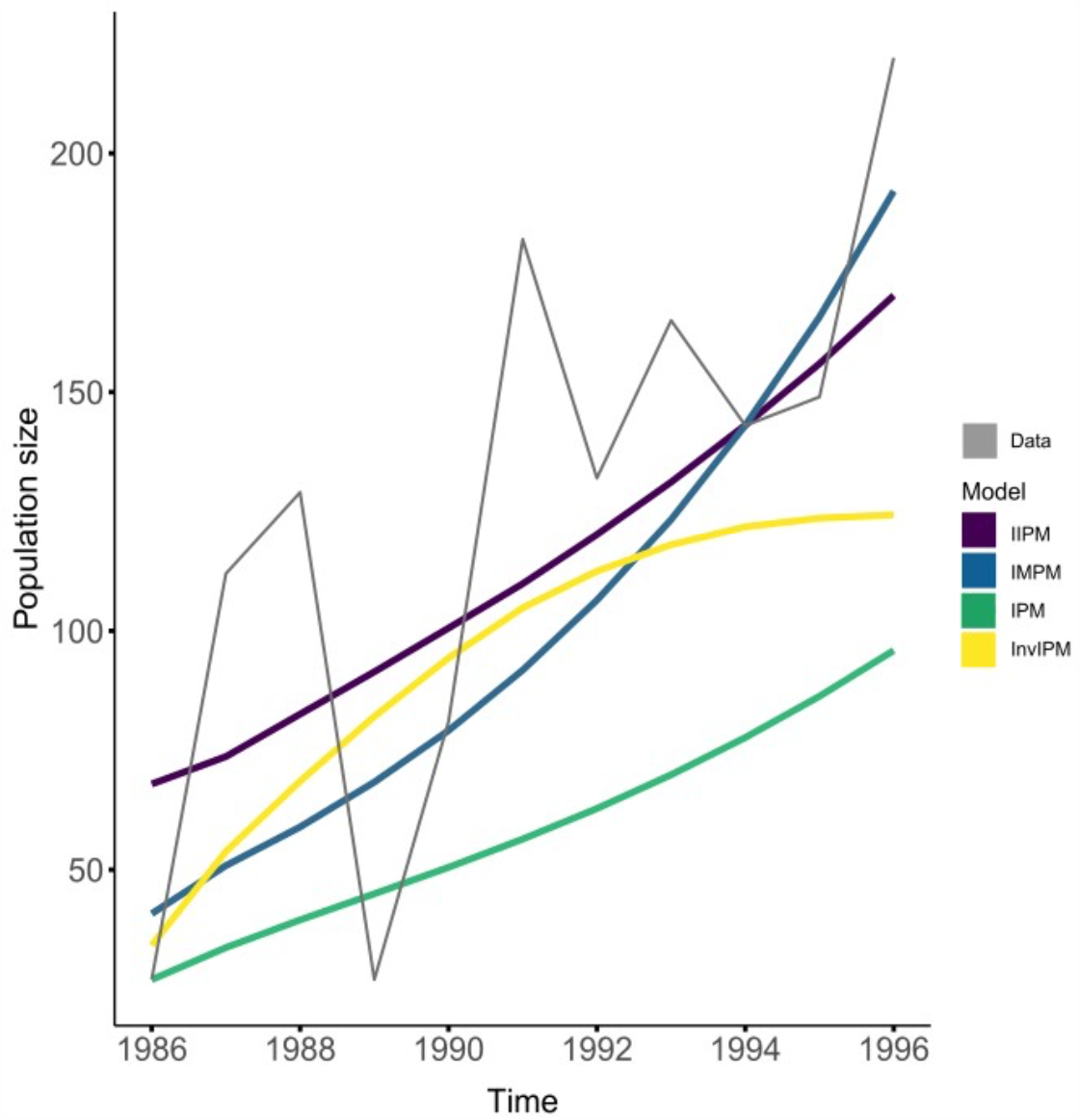
Predicted sizes of the Soay sheep population over 11 years according to the four population models compared in this study: IIPM: Integrated Integral Population Model, IPM: Integral Population Model (IPM), Inverse Integral Population Model (InvIPM), IMPM: Integrated Matrix Population Model. The observed population size time series is in grey.

## Discussion

Matrix population models existed for almost six decades (Leslie, 1945) before the population was structured around a continuous variable giving birth to the so-called integral projection models (Easterling et al., 2000). Although IPMs are mostly used for plant populations, their application in animal demography has increased over time (Merow et al., 2014; Salguero-Gómez et al., 2015; 2016). Two years after the proposal of the IPM, the idea of incorporating information from the population level into demographic models emerged in animal demography context with the integrated (matrix) population models (Besbeas et al., 2002). Although it would seem that the logical next step would have been the close emergence of the integrated version of the IPM, a delay of 20 years may be largely due to a gulf of communication between animal and plant demographers.

Here, we close this existing gap by presenting the version of the IMPM when the state variable is continuous: the Integrated Integral Population Model (IIPM). This model emerges from the logical argument behind IPMs and InvIPMs. From the former models, the IIPM inherits the inclusion of vital rates information determining the dynamics of a population (Easterling et al., 2000; Merow et al., 2014; Ellner et al., 2016). From the latter model, the IIPM inherits the capacity to describe observed population trends (González et al., 2016).

The IIPM reconstructs population dynamics taking into account information from both the individual and population levels. This is achieved by placing the IPM structure (eq. 1) at the core of the statistical fitting of the individual and population data. Given that the IPM links the individual processes (survival, growth, and reproduction) with the population emerging properties (size and structure), incorporating the IPM structure within the statistical fitting of the IIPM creates feedback that constrains them both to be consistent among each other. By doing so the IIPM solves the input-output mismatch problem of IPMs and InvIPMs. Our results validate this assertion: the IPM produced the best estimates at the individual level and the worst at the population level (Table 1), while the InvIPM performed in the opposite way but, as it is clear from Fig. 2, having only population level data can produce very distorted vital rates. In turn, the IIPM produced sensible results both at the individual and population levels (Figs 2-4.). Thus, the IIPM inherits the capacity of each model to correctly describe the data they use but avoids the lack of fit that they display when confronted with data at the alternative organisational level.

### IIPMs & IMPMs

IIPMs are the complement to IMPMs, as they provide a tool to model in an integrated manner the dynamics of a structured population when the structuring variable is of a continuous nature. By combining different sources of demographic data into one statistical model, in our case survival, reproduction, population sizes and structures, IMPMs can increase the precision of our demographic parameters and even estimate parameters for which data is not available (Schaub et al., 2007; Schaub & Abadi, 2011). As demonstrated by our results, IMPMs and IIPMs follow the same idea of placing the population model at the core of the statistical fitting procedure. By doing so, they can produce parameters estimates that reflect all the information available on a species population dynamics.

IMPMs emerged in the context of animal studies where there was a need to fit population models using fragmentary demographic data (Besbeas et al., 2002; Schaub et al., 2007), and later it was realised that they allow to model both individual and population level data (Schaub & Abadi, 2011). In our case, we saw the IIPM as the combination of the IPM and the InvIPM. Nonetheless, the result is the same: integrated population models, i.e., IMPMs and the IIPMs, allow the estimation of both vital rates and population attributes, and these estimates are more precise that those provided by their non-integrated counterparts.

### Disadvantages of IIPMs

The IIPM reconstructs multilevel population dynamics at the expense of a greater programming effort when compared with non-integrated models. This explains, at least partially, the reason why integrated models have had relatively limited applications in comparison with non-integrated ones.

We developed the IIPM in Stan (Stan Development Team, 2023) because it is a flexible software that allows the inclusion of as many model details as required. However, this flexibility requires learning Stan’s particular language, how to code the IPM structure in this language and explicitly formulate the likelihoods associated with the vital rates and population attributes. Although we use Stan, the IIPM can be coded into other similar languages such as NIMBLE (NIMBLE Development Team, 2016), JAGS (Plummer, 2003) and WinBugs (Kéry & Shaub, 2011); actually, existing IMPMs have been implemented in NIMBLE (Gamelon et al., 2021) and JAGS (Riecke et al., 2019), and the IPM^2^ in NIMBLE (Plard et al., 2019b).

### IPM^2^ vs IIPM

Recently, Plard et al. (2019b) presented the IPM^2^. Counterintuitively to its name, this model seeks to combine an MPM with an IPM by dividing the population into two discrete stages (juvenile and adults), and then relating, within each stage, the vital rates with a continuous state variable (laying date of the first annual brood in a population of barn swallows, *Hirundo rustica*). Rather than an IIPM, this is an integrated hybrid model that combines an MPM with an IPM by having a discrete state variable and a continuous one structuring the population (Ellner et al., 2021). This hybrid model goes beyond the traditional structured population models in which a population is structured by a single state variable. Our objective was a more humble one: to present the continuous version of an IMPM, emphasising the differences between our IIPM and closely related models: the IPM, the InvIPM and the IMPM.

### Integrated Integral Population Model: why the name?

*Population* is the term most often associated with models describing the dynamics of populations. A quick survey of titles containing the term *population* in combination with the terms *model* and *matrix/integral* yielded over 10,000 more articles than a survey of titles containing the term *projection* combined with these terms. This is mainly because matrix population models have been constructed for almost 80 years (Leslie, 1945) and integrated (matrix) population models for over 20 (Besbeas et al., 2002).

*Projection* was the first term used to name integral projection models (Easterling et al., 2000). This term emphasises the iterative nature of structured population models; however, we must recognize that iteration is not a common feature of IPM applications. Actually, most IPM applications focus only on the asymptotic attributes of the kernel such as the asymptotic population growth rate, its sensitivities and elasticities, stable population structure and reproductive values (Crone et al., 2011; Kendall et al., 2019), which are computed directly from the kernel. These attributes allow us to analyse and make decisions in many areas of biology such as species conservation (Crone et al., 2011; Lown et al., 2020), invasive species control (Ramula et al., 2010) and species management (Schmidt et al., 2011; Isaza et al., 2017). However, we need to recognize that 1) these attributes will be biassed if they come from non-integrated models; 2) the dynamics of natural populations are never static and studying asymptotic attributes can have low informative value, and 3) focusing only on this core asymptotic properties hides the low capacity that IPMs can have in describing population attributes.

Therefore, as we endeavour to accentuate the importance of the population attributes in the fitting of a population model and to use the most commonly used term among demographers, we decided to prioritise the term *population* rather than *projection* in naming our model.

### Further complications

Here, we used the simplest structure of the IIPM to exemplify it. However, its structure can be easily modified to take into account intrinsic and extrinsic factors that may affect a population’s dynamics as well as methodological issues that may have an impact on the estimation of individual-or population-level parameters. Intrinsic factors include all potential variables, inherent to the individual (e.g., size, sex, and age), that may produce fitness variation among individuals (Riecke et al., 2019; Ellner et al., 2016; Fung et al., 2022); we can even include the individual as a random effect to address dependencies among databases sharing the same individuals (Ellner et al., 2016; Fung et al., 2022). Extrinsic factors include both abiotic and biotic environmental variables (e.g., climate, intraspecific interactions) that may also affect an individual’s fitness (Ellner et al., 2016; Plard et al., 2019a; 2019b). These variables can be included as explanatory variables in the vital rates regression structure. This structure varies on the number and type of state and environmental variables and depends entirely on the life history of each species and the research questions we want to address, which can go beyond population demography such as eco-evolutionary dynamics and the forecast of demographic behaviour of species facing climate change (Vindenes & Langangen, 2015; Ellner et al., 2016; Plard et al., 2019a; 2019b).

Methodological issues include, but are not restricted to, quantification errors. These errors can occur both at the individual and population levels. At the individual level, IIPMs can incorporate imperfect detectability and marker loss rates (Gibson et al., 2013; Riecke et al., 2019). At the population level, IIPMs can incorporate population observation error (Shaub & Abadi, 2007; Besbeas & Morgan, 2019). These quantification errors can also be included in the IIPM as random effects and their inclusion in the model helps in the fitting procedure (Tavecchia et al., 2009; Ellner et al., 2016). Nevertheless, integration does not solve other methodological issues such as model misspecification, which may have important effects on IPM-derived metrics (González et al., 2013).

### Implications for the monitoring of species dynamics

We can take advantage of the fact that the IIPM is fed by individual and population data to optimise the sampling design for population dynamics studies. As IPMs and MPMs are fed only by individual data, their sampling designs focus on obtaining vital rates data for each individual. Thus, it is necessary to mark and follow each individual for at least two years (Merow et al., 2014; Ellner et al., 2016). In the same way, as InvIPMs are fed only by population data their sampling design focuses do not require individual marks. This allows us to avoid sampling issues such as marker loss and vital rates measurement errors (Gross et al., 2002; Ghosh et al., 2012). In any of these non-integrated models, a small sample can lead to a misconstructed population dynamics (González et al., 2016). Frequently, small samples are more common at the individual level than at the population level. In the IIPM these individual small samples can be compensated by population samples. This compensation is based on the fact that the year-to-year variation in the times series of population attributes reflects the set of individual performances (Gross et al., 2002; Ghosh et al., 2002; 2012; Ellner et al., 2016). Thus, if we use an IIPM, it may be sufficient to sample the most influential vital rates, e.g., those with higher elasticities, from a small sample of individuals and allocate a larger sampling effort to quantify population attributes, which are inherently easier to obtain.

## Conclusion

The objective of a structured population model should be to correctly describe both individual population patterns. Non-integrated population models, such as IPMs, MPMs and InvIPMs, by not using individual and population data simultaneously, fail in this objective. We propose the IIPM which is a continuously structured population dynamics model that places the entire population model at the core of the statistical fitting procedure. By integrating the logic behind the IPM, InvIPM and IMPM, the IIPM inherits their benefits: the fit of individual data of the IPM, the fit of population data of the InvIPM, and the capacity to incorporate different information sources into a single model of the IMPM. Therefore, IIPM reaches the sought objective.

## Acknowledgements

We thank Carlos Martorell and Alberto Búrquez for their helpful comments on previous versions of this manuscript. This paper serves as a fulfilment of PP-T to obtain her master’s degree in Biological Sciences (Ecology) in the Posgrado en Ciencias Biológicas, UNAM. We thank CONAHCYT for proving a scholarship to PP-T.

